# Free-living food intake and repopulation of the gut microbiota after a health-screening colonoscopy

**DOI:** 10.1101/2025.02.26.640308

**Authors:** Yezaz A Ghouri, Aaron C Ericsson, Jennifer M Anderson, Jessica G George, Elizabeth J Parks, Katherene OB Anguah

## Abstract

**Objective:** Role of microbiome has been highly studied for its association with various medical conditions. After a colonoscopy, repopulation of colonic microbial load is known to occur, however the quality and timing of natural repopulation has not been investigated after a bowel preparation. Further, no study has documented detailed free-living dietary intakes concurrently with gut microbiome repopulation post-colonoscopy. Here we sought to determine the early pattern of repopulation relative to dietary intake.

**Methods:** Healthy adults (n=15 [4 female/11 male], BMI=27.2±3.9 kg/m^2^, age 51.4±7.2 y) who were scheduled to undergo a screening colonoscopy were recruited from the Gastroenterology Clinic at the University of Missouri. Within two weeks before the colonoscopy (baseline), subjects completed detailed food records for 3 days. Post-colonoscopy, subjects ate their free-living diets and detailed food records were collected on Days 0, 1, 2, 4, 7, 10, and 13. Fecal samples were obtained pre-colonoscopy and on post-colonoscopy Days 3, 5, 8, 11, and 14. Gut microbiome composition was assessed by 16S rRNA amplicon sequencing.

**Results:** Within 5 days after the procedure, subjects reported consuming more total daily energy relative to baseline, presumably to make up for the low energy intake that occurred during the bowel-prep. At baseline, fiber intake (21.0±9.1 g/d) was higher than on the day of the colonoscopy, Day 0 (16.1±11.2, *P*=0.0159). Thereafter, daily fiber intake was the same as baseline. Marked intersubject microbiome beta diversity was observed by principal coordinate analysis using weighted and unweighted dissimilarities (*P*=0.0001, F=15.23, one-way PERMANOVA). Select taxa were depleted acutely post-colonoscopy (e.g., within the phylum Bacillota). Specifically, significant effects of time were observed between baseline and Day 3 fecal samples (pairwise *P*=0.0013, F=2.9). These changes trended to return to baseline by Day 5 and with subsequent samples, taxa remained similar to baseline when tested using a weighted dissimilarity analysis (Bray-Curtis).

**Conclusions:** These results quantitatively demonstrate the magnitude of the significant changes in microbial relative abundance and diversity immediately post-colonoscopy. The timing of repopulation aligned with changes in fiber intake after the procedure. These data highlight the importance of nutrition after a screening colonoscopy in reestablishing a healthy microbiome.

## Introduction

Within the last two decades, the contribution to human health of the gut microbiome has highlighted its role in modulating the risk for certain chronic conditions such as obesity, type 2 diabetes, inflammatory bowel disease, and cardiovascular disease(1, 2)]. Colonoscopy is a routine procedure performed in the U.S. to screen for colorectal cancer, and every year, over 15 million people in the U.S. undergo screening colonoscopies(3)]. Due to the substantial rise in cases of early onset colorectal cancer, the recent recommendation from the American Cancer Society is for screening to begin at 45 years of age for people with average risk for colorectal cancer regardless of race(4)]. To prepare for colonoscopy, a standardized colon preparation is designed to cleanse the bowel of its stool contents which significantly reduces the microbial population. This is achieved with the use of laxative consumed in a large volume. Yet, little is known about how the gut microbiome repopulates after this procedure.

Four publications have described the changes in the microbiome post-colonoscopy. These studies demonstrated a 31-fold reduction in microbial load(5)] or a general reduction in microbiota composition(6, 7)] and diversity (8)] immediately following the colonoscopy procedure which returned to baseline levels within 2 weeks post-colonoscopy. Thus, standardized screening colonoscopy procedures are likely significantly reducing the microbiome. However, the pattern of repopulation of microbes after the procedure has not been clearly documented in individuals. Further, given the importance of diet in affecting the pattern of microbial species (9), to understand the repopulation process, dietary intake should be documented. No study to date has collected detailed food intake records under free-living conditions in individuals before or immediately after the colonoscopy procedure. The goal of the present study was to determine the early pattern of microbial repopulation relative to diet in healthy subjects before undergoing colon preparation and over a period of two weeks after their colonoscopy procedure.

## Materials and methods

### Study participants

Fifteen patients scheduled to undergo health-screening colonoscopies were recruited from the Missouri Digestive Health Center at the University of Missouri (MU) – Columbia. Subjects were consented and enrolled sequentially from July through October 2023. Inclusion criteria were men and women, aged 45-65 years, without active GI disease (e.g., colon cancer, active GI bleeding, inflammatory bowel disease etc.) and willing to provide fecal samples. Subjects were excluded if they 1) were on medications known to affect the gut microbiome (e.g. antibiotics within the past 30 days), appetite, and/or body weight, 2) had a history of GI disease or 3) lived more than 45 miles away from the University of Missouri, as a result of the requirement that fecal samples be delivered six times over a two week period. This study was approved by the MU Institutional Review Board (Protocol #2053546) and registered (clinicaltrials.gov NCT06023940). Of note, this study has multiple arms, all data from the current analysis come from data that were collected after clinical trial registration for that arm and are observational in nature (no intervention given). The first subject enrollment (signed consent) for the study arm whose data are reported in this manuscript began on July 10, 2023 and the last subject completed the study on October 30, 2023. All subjects enrolled in the study gave written informed consent with HIPAA authorization.

As shown in **Fig 1**, 157 participants were initially contacted to determine eligibility. Out of this, 25 were eligible and scheduled for an in-person screening visit, with 19 participants completing the informed consent process. Four out of these 19 subjects were excluded after the informed consent process, resulting in 15 subjects that completed the study.

**Fig 1.**
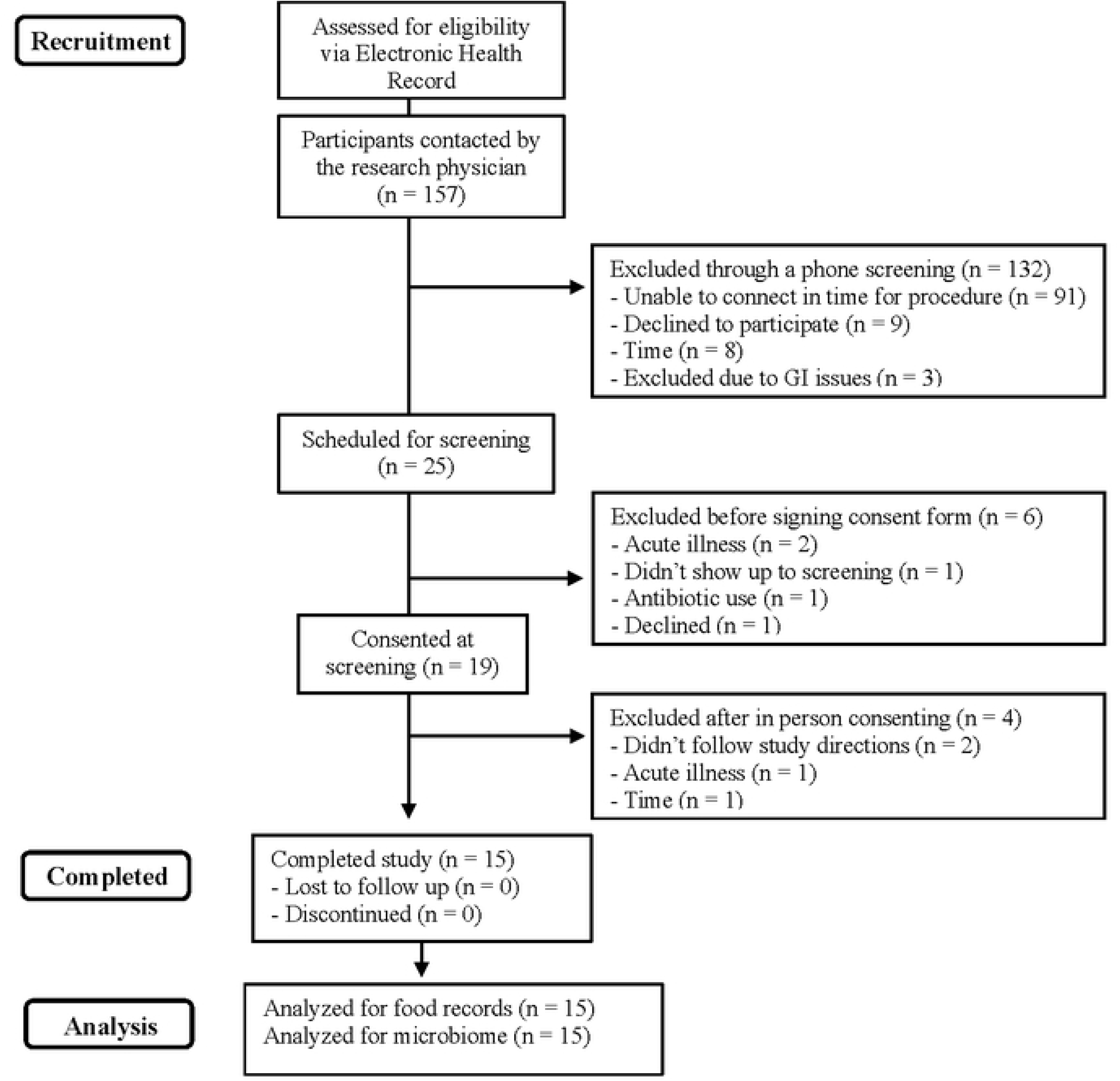
Consort flow diagram.

### Study design

The study design is shown in **Fig 2**. Prospective patients were identified by screening the upcoming colonoscopy appointments. Subsequent electronic health record charts were reviewed to determine potential eligibility. If subjects were preliminarily eligible and interested in participating in the study, written informed consent was obtained during an in-person screening visit prior to their scheduled colonoscopy. Eating behavior was assessed using the Three-Factor Eating Questionnaire (TFEQ-51-item questionnaire with three subscales: a 21-item dietary restraint scale, a 16-item dietary disinhibition scale, and a 14-item hunger scale) which was originally developed by Stunkard and Messick (10). The TFEQ was administered to subjects at the screening visit in a fasted state.

**Fig 2.**
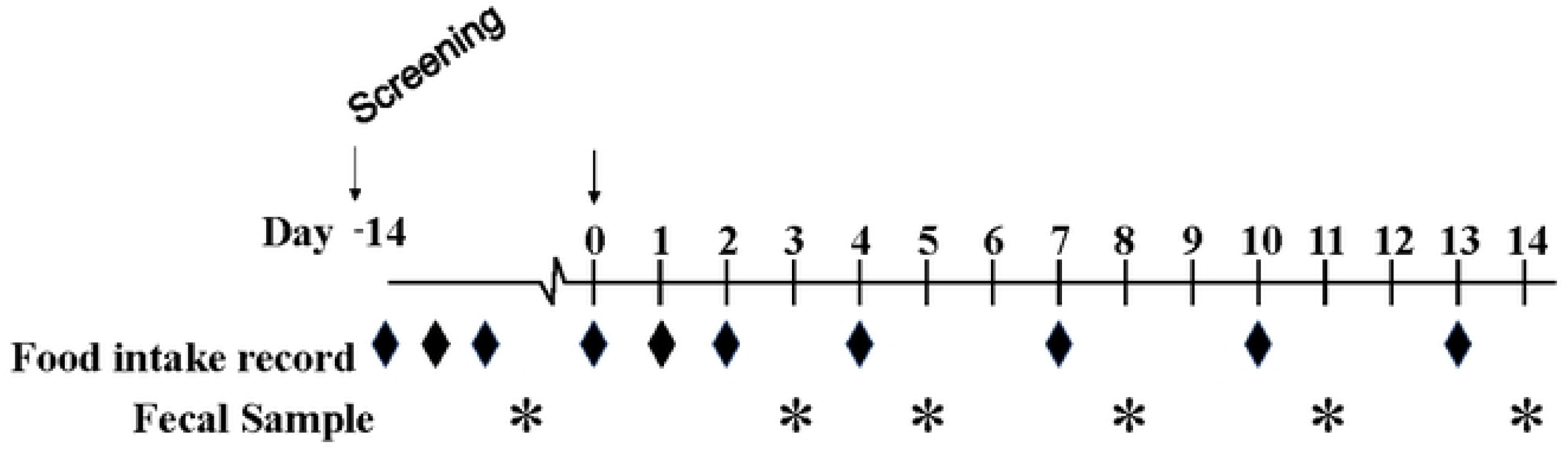
Study design. Subjects completed food records and provided fecal samples before and after their health-screening colonoscopies.

Enrolled subjects completed ten days of food records including a 3-day food record completed after the in-person screening visit (within two weeks pre-colonoscopy) and on seven days following the colonoscopy: Days 0 (procedure day), 1, 2, 4, 7, 10, and 13. Subjects also collected fecal samples at baseline (1 sample within two weeks pre-colonoscopy) and after the procedure on Days 3, 5, 8, 11, and 14. In addition, they also completed the Bristol Stool Chart for every fecal sample collected. Subjects did not collect food record data or fecal samples the day before their medical procedure, as this was the day of their colonoscopy preparation. Per the standard directions from the healthcare team, patients followed a clear liquid diet the day before the procedure and ingested 3.8 L of polyethylene glycol 3350 (GOLYTELY^®^) split into two doses, 1.9 L administered the night before the procedure and 1.9 L the morning of the procedure. The subjects were n.p.o. for at least 2 hours before the procedure. All the patients underwent sedation with propofol for the procedure. Following the procedure, diet was resumed when they were able to tolerate oral intake.

### Blood biochemistries

Blood biochemistries data were collected from the electronic health record. The most recent data, back to 12 months prior to the colonoscopy were included. Not all subjects had blood biochemistry data available within 12 months of their procedure.

Participants medical health records screening start date was June 12, 2023, and end date was October 3, 2023. None of the authors had access to information that could identify individual participants during or after data collection, except one author (Anderson JM) who conducted the medical records screening. However, data were de-identified after collection.

### Dietary intake

With regard to dietary intake, subjects were instructed to consume their usual diet during the study. Instructions on how to fill out the food records were provided by the study dietitian, who reviewed the records and interviewed the subject to verify and document any missing details. Dietary data were entered in Nutrition Data System for Research® 2021 (Nutrition Coordinating Center, University of Minnesota) to determine nutrient quantity and composition and then the data were analyzed in Excel®.

### DNA isolation, and quantification

The stool DNA was extracted using PowerFecal kits (Qiagen) according to the manufacturer instructions, with the exception that samples were homogenized in the provided bead tubes using a TissueLyser II (Qiagen, Venlo, Netherlands) for ten minutes at 30 Hz (cycles/sec), rather than performing the initial homogenization of samples using the vortex adapter described in the protocol, before proceeding according to the protocol and eluting in 100 µL of elution buffer (Qiagen). DNA yields were quantified via fluorometry (Qubit 2.0, Invitrogen, Carlsbad, CA) using quant-iT BR dsDNA reagent kits (Invitrogen) and normalized to a uniform concentration and volume.

### 16S rRNA library preparation and sequencing

Library preparation and sequencing were performed at the MU Genomics Technology Core. Bacterial 16S rRNA amplicons were constructed via amplification of the V4 region of the 16S rRNA gene with universal primers (U515F/806R) previously developed against the V4 region, flanked by Illumina standard adapter sequences (11, 12)]. Dual-indexed forward and reverse primers were used in all reactions. PCR was performed in 50 µL reactions containing 100 ng metagenomic DNA, primers (0.2 µM each), dNTPs (200 µM each), and Phusion high-fidelity DNA polymerase (1U, Thermo Fisher). Amplification parameters were 98°C^(3 min)^ + [98°C^(15 sec)^ + 50°C^(30 sec)^ + 72°C^(30 sec)^] × 25 cycles + 72°C^(7 min)^. Amplicon pools (5 µL/reaction) were combined, thoroughly mixed, and then purified by addition of Axygen Axyprep MagPCR clean-up beads to an equal volume of 50 µL of amplicons and incubated for 15 minutes at room temperature. Products were then washed multiple times with 80% ethanol and the dried pellet was resuspended in 32.5 µL EB buffer (Qiagen), incubated for two minutes at room temperature, and then placed on the magnetic stand for five minutes. The final amplicon pool was evaluated using the Advanced Analytical Fragment Analyzer automated electrophoresis system, quantified using quant-iT HS dsDNA reagent kits, and diluted according to Illumina’s standard protocol for sequencing as 2×250 bp paired-end reads on the MiSeq instrument.

### Informatics analysis

The DNA sequences were assembled and annotated at the MU Informatics Research Core Facility. Primers were designed to match the 5’ ends of the forward and reverse reads. Cutadapt (13) (version 2.6) was used to remove the primer from the 5’ end of the forward read. If found, the reverse complement of the primer to the reverse read was then removed from the forward read as were all bases downstream. Thus, a forward read could be trimmed at both ends if the insert was shorter than the amplicon length. The same approach was used on the reverse read, but with the primers in the opposite roles. Read pairs were rejected if one read or the other did not match a 5’ primer, and an error-rate of 0.1 was allowed. Two passes were made over each read to ensure removal of the second primer. A minimal overlap of three bp with the 3’ end of the primer sequence was required for removal. The QIIME2(14)] DADA2(15)] plugin (version 1.10.0) was used to de-noise, de-replicate, and count amplicon sequence variants (ASVs), incorporating the following parameters: 1) forward and reverse reads were truncated to 150 bases, 2) forward and reverse reads with number of expected errors higher than 2.0 were discarded, and 3) Chimeras were detected using the “consensus” method and removed. R version 3.5.1 and Biom version 2.1.7 were used in QIIME2 to analyze the samples. Taxonomies were assigned to final sequences using the Silva.v132(16)] database, using the classify-sklearn procedure.

### Statistical analysis

Microsoft Excel and Statistical Package for Social Sciences (SPSS) version 29.0.1.0 (171) were used to analyze the subject characteristics and dietary data. All baseline continuous variables were expressed as mean ±SD together with the range, while categorical variables were expressed as n or %. For the dietary data, each participant’s total energy intakes (kcal/day) averaged for the 3-days within the two weeks prior to the colonoscopy procedure. These baseline values were set at 100% intake and the intakes after colonoscopy were expressed as a percentage relative to this baseline. Statistical analysis for the microbiome data was conducted using Paleontological Statistics Software Package (PAST, University of Oslo, Norway).

Beta diversity analyses to compare differences in microbial composition between different subjects, and across different time points, were conducted using Principal Component Analysis (PCA). Multivariate analysis, specifically, one-way PERMANOVA analysis was conducted to test differences in composition, using both unweighted similarity (Jaccard) and weighted similarity (Bray-Curtis) measures. Pairwise comparisons were used to compare results from different timepoints. The rarefied data were used in the microbiome statistical analyses. Two subjects had one each of total ASV coverage for one timepoint being below the threshold counts per sample (one subject at Day 3 (D3) and a second subject at D11). Thus, these two timepoints were not included in the microbiome analyses, leaving data from 4 instead of 5 timepoints for these two subjects. Wilcoxon signed rank tests comparing relative abundance of all ASVs between D0 and D3, were conducted and only ASVs that yielded a raw *P* value < 0.05 were reported. Furthermore, correction for multiple tests was conducted using false discovery rate (FDR) test.

Pearson’s correlational analyses were run to investigate the associations between dietary intakes and the relative abundance of the first 10 ASVs with the least *P*-values. Due to the exploratory nature of these analyses, we did not correct the *P*-value for multiple comparisons. Differences were considered statistically significant at *P*≤0.05 in all analyses conducted.

## Results

### Participant characteristics and endoscopy results

Study participants were predominantly white males (73.3%) with an average BMI in the overweight category and ranging from 20.5-33.9 kg/m^2^ (**Table 1**). Blood biochemistry results were generally within a healthy range. The average Bristol Stool Chart rating at the screening visit was 3±1. At this time, 66.7% of the participants rated their fecal samples in the typical range (Type 3, 4, or 5). On D3 and D14 post-colonoscopy, 86.6% and 73.3% respectively of the participants rated their fecal sample as Type 3, 4, or 5. With regards to eating behavior, on average subjects’ had low restraint, disinhibition, and hunger scores.

**Table 1.**
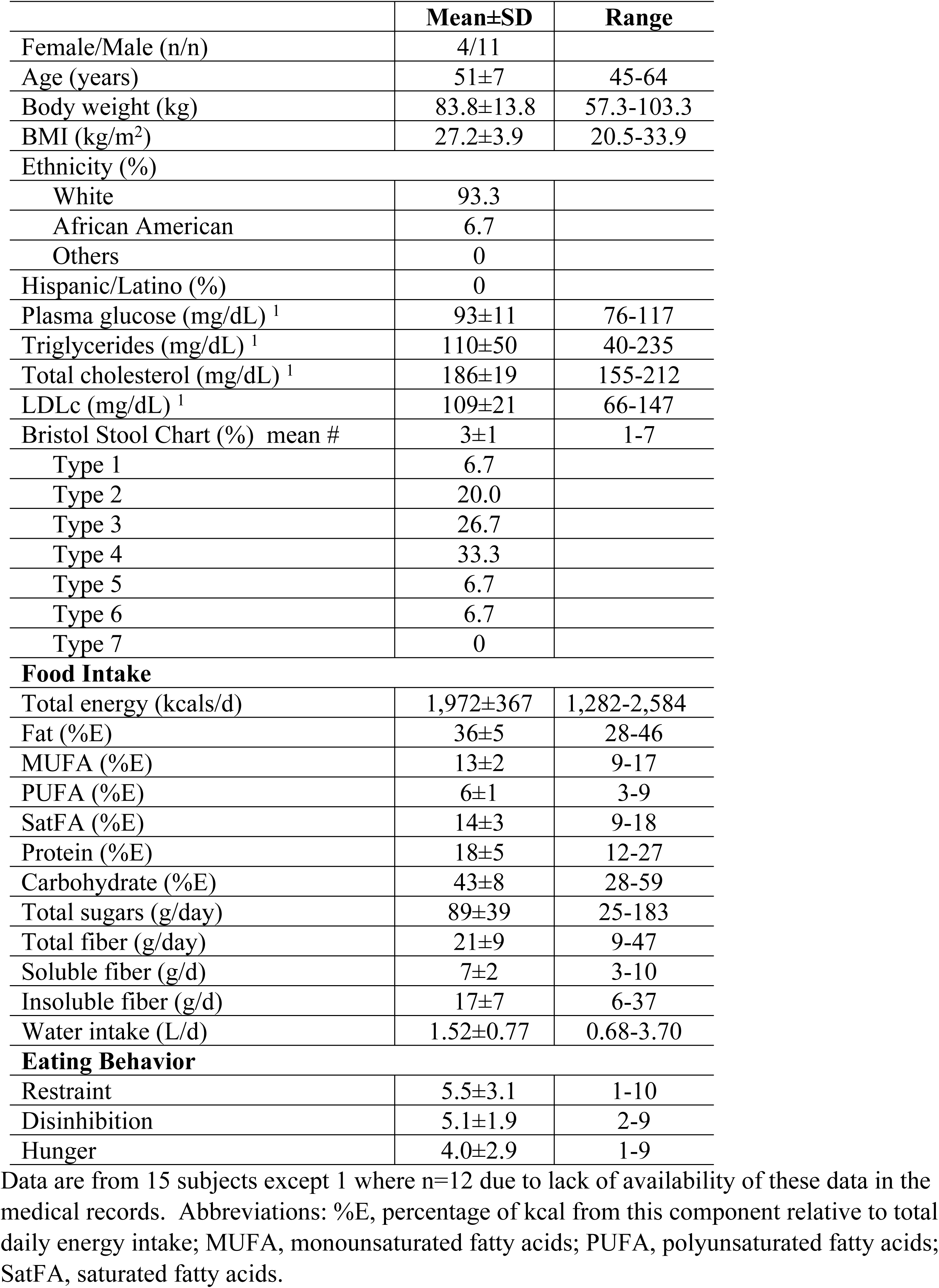
Participant characteristics and food intake at baseline.

Polyps were found and removed during the colonoscopy procedure in six of the fifteen study participants, which was not an unexpected result. Five of the six participants had ≤2 polyps, while one participant had 7 polyps.

### Dietary intake

With regard to food intake (T**able 1**), at baseline prior to the colonoscopy procedure, study participants consumed 36±5 % fat, 18±5% protein, 43±8% carbohydrates, which is generally more protein and less carbohydrate as a percentage of energy compared to the average American adult(17)]. Furthermore, participants ate more fiber than the average American (21 g/day vs ∼15 g/day)(17)], and ate within the recommended limit of added sugars (<50 g/day for 2000 kcal/day)(18)] (**Table 1**). Baseline fiber intake (21.0±9.1 g/day) was higher than on the day of the colonoscopy, Day 0 (D0) (16.1±11.1, *P*=0.0159). Subsequently, daily fiber intake was the same as baseline. As shown in **Fig 3**, participants had a variable energy intake throughout the study. During the 2 weeks post-colonoscopy, all subjects reported consuming more than the pre-colonoscopy (baseline) intake; this increased intake was likely to compensate for the lack of food intake on the day before the colonoscopy. Most subjects exhibited a peak total energy intake occurring before Day 5 (D5), although three subjects (T017, T020, T038) also reported peaks in energy intake later in the observation period.

**Fig 3.**
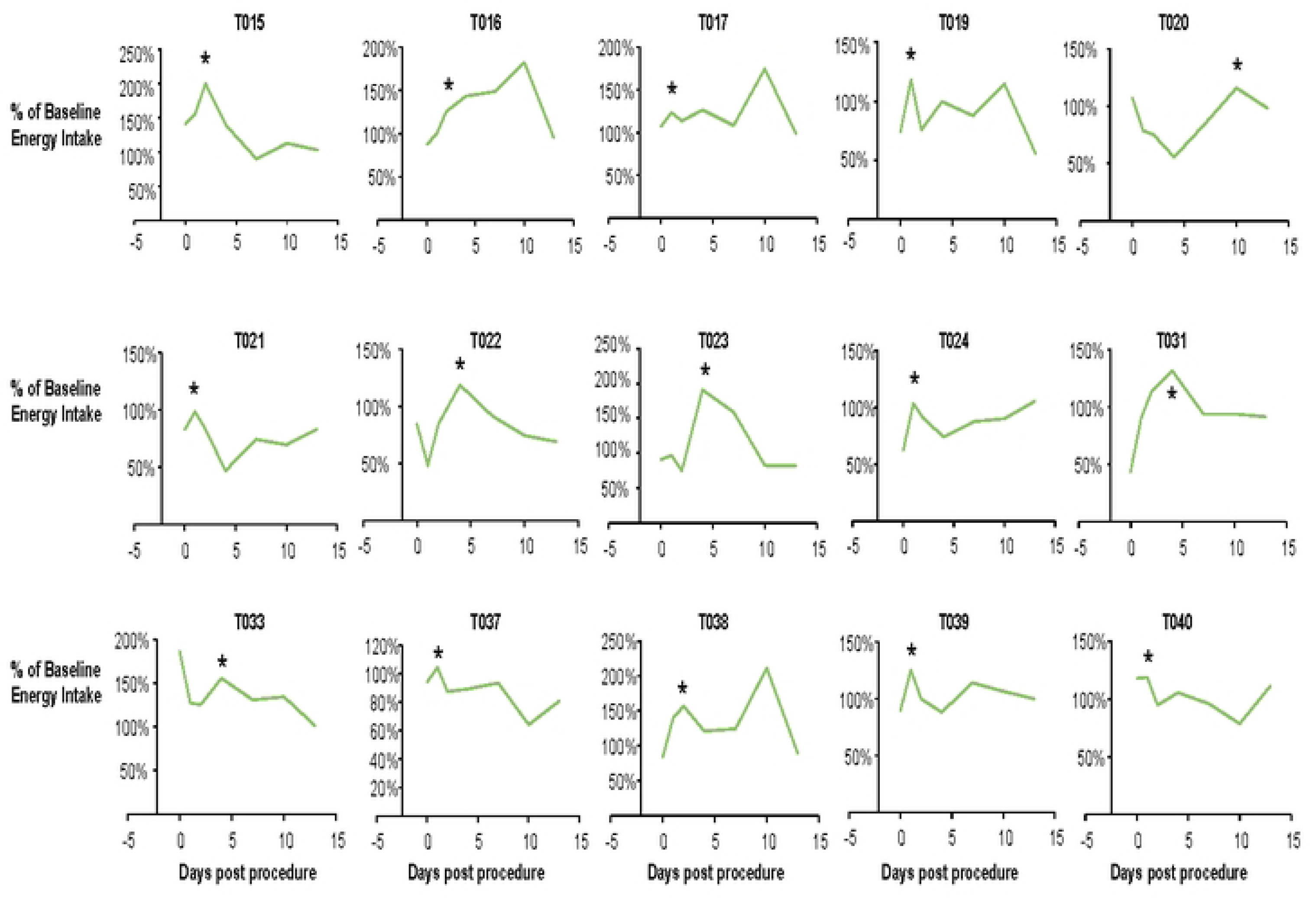
Changes in food intake after the colonoscopy relative to pre-colonoscopy intake. Each figure represents data from a single subject. Food intake (kcals/day) on each of the post-colonoscopy days is presented relative to the baseline, free-living intake that was assessed over three days in the two weeks before the procedure. * Denotes the timing of an early observed peak in food intake after the colonoscopy.

### Microbiome results

Beta diversity assessment using principal coordinate analysis indicated significant intersubject beta diversity using weighted and unweighted dissimilarities (*P*=0.0001, F=15.23, one-way PERMANOVER) (**Fig 4A**). **Fig 4B and 4C** show beta-diversity results for time effects. While the overall effect of time was modest, pairwise comparisons revealed significant effects of time between pre-colonoscopy and Day 3 post-colonoscopy (D3) samples (*P*=0.0015, F=2.91), however there were no significant effects of time between pre-colonoscopy and any other time point beyond D3 post-colonoscopy. These differences were observed using a weighted dissimilarity (Bray-Curtis-Fig 3C), but not with Jaccard dissimilarity (**Fig 4B**). Furthermore, D3 post-colonoscopy samples were not significantly different from D5 samples, and the difference between D5 and the remaining timepoints diminished gradually over time (**Fig 4C**). These data support the concept that the microbiome had achieved a new steady state of composition.

**Fig 4.**
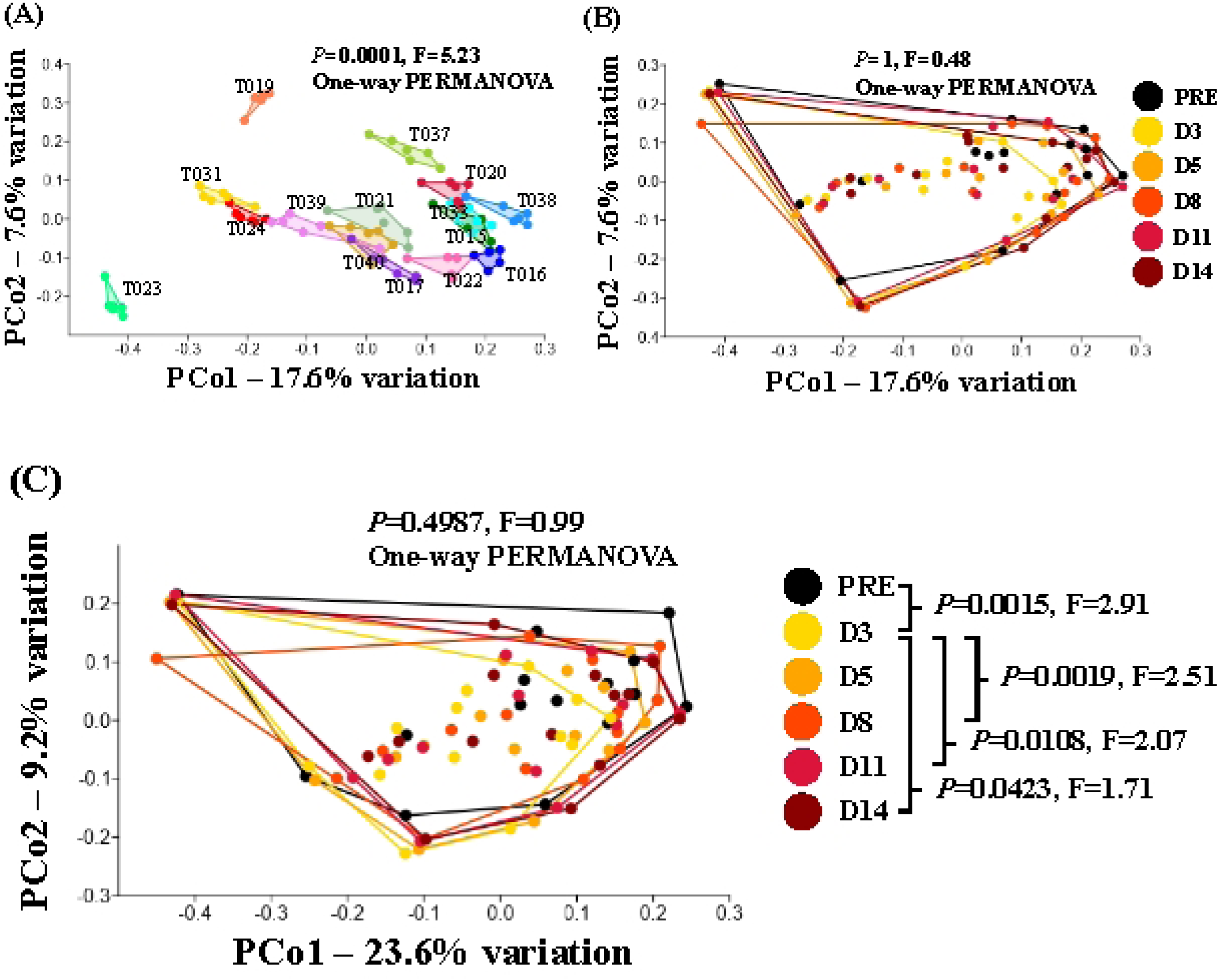
Principal coordinates analysis. **(A) Inter-subject beta diversity using Jaccard dissimilarities.** T0XX represents individual subject identifiers. **(B)** Beta diversity over time using Jaccard dissimilarities, and **(C)** Beta diversity over time using Bray-Curtis dissimilarities. PRE, denotes the pre-colonoscopy data, and for post colonoscopy samples: D3 is Day 3, D5 is Day 5, D8 is Day 8, D11 is Day 11, D14 is Day 14.

Because pre-colonoscopy samples were only different from D3 samples using the weighted dissimilarity comparison, we performed serial Wilcoxon rank tests on all ASVs to identify taxa contributing to the observed change in beta-diversity. While 49 ASVs yielded a raw *P* value<0.05, none withstood correction to control the false discovery rate. Collectively however, these taxa are interpreted to represent the subtle changes in beta-diversity observed at D3 post-colonoscopy. A heat map was generated to visualize the data from these two timepoints. As can be seen in the top 75% of the heat map (**Fig 5**), select taxa (primarily *Bacillota*) appeared to be depleted immediately post-colonoscopy, while the lower 25% of the heat map shows a handful of ASVs that were differentially enriched post-colonoscopy. Example ASVs that were reduced from baseline in relative abundance at D3 included the genera [Clostridium]_methylpentosum_group, Clostridia_vadinBB60_group, UCG-010, *Roseburia*, uncultured, *Intestimonas*, Clostridia_UCG-014, *Ruminococcus*, *Colidextribacter*, *Medibacter*, and Christensenellaceae_R-7_group etc. all belonging to the phylum Bacillota. The few ASVs in the bottom 25% of the heat map that were differentially enriched at D3 included the genera *Parabacteroides* and *Bacteroides,* both belonging to the phylum *Bacteroidota*; the genera *Lachnoclostridium*, *Veillonella*, and *Gemella* belonging to the phylum Bacillota; and an unresolved genus in the family *Pasteurellaceae*, phylum *Pseudomonadota*. Shown in **Fig 6**, are violin plots indicating the first 4 ASVs with the lowest raw *P*-values as an example of the marked differences between pre-colonoscopy and D3 samples.

**Fig 5.**
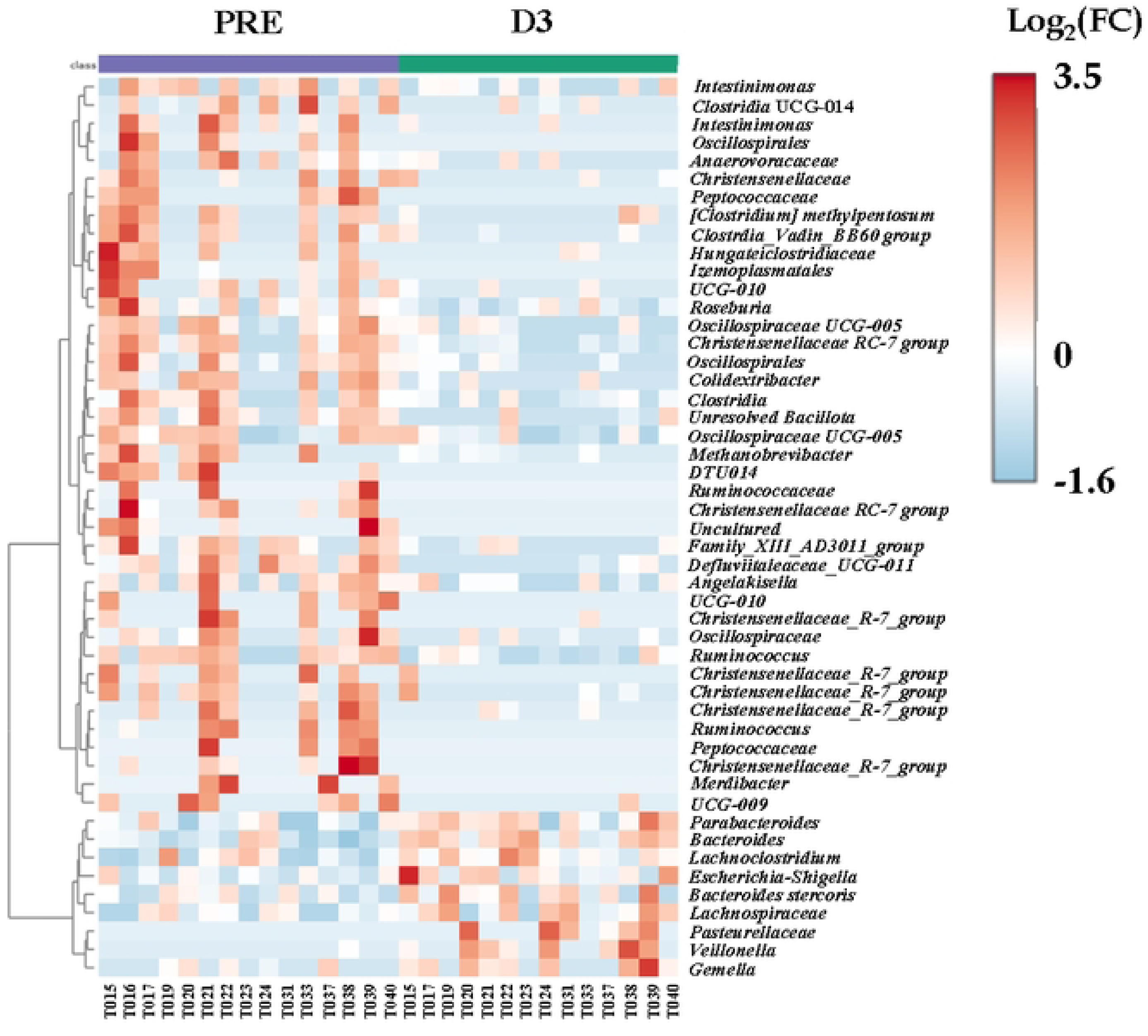
Heat map showing differentially abundant ASVs at the Genus level between pre-colonoscopy and Day 3 post-colonoscopy (D3) samples. The color of the boxes corresponds to the normalized and log-transformed relative abundance of the ASVs. The first half of the x-axis represents individual subject data during pre-colonoscopy(40), while the second half represents the same subjects at Day 3 post-colonoscopy (D3). One subject (T016) was missing Day 3 datum after the raw data were rarefied due to having very low microbial count. For ASVs with unresolved genus classification, the next available taxonomic classification was used.

**Fig 6.**
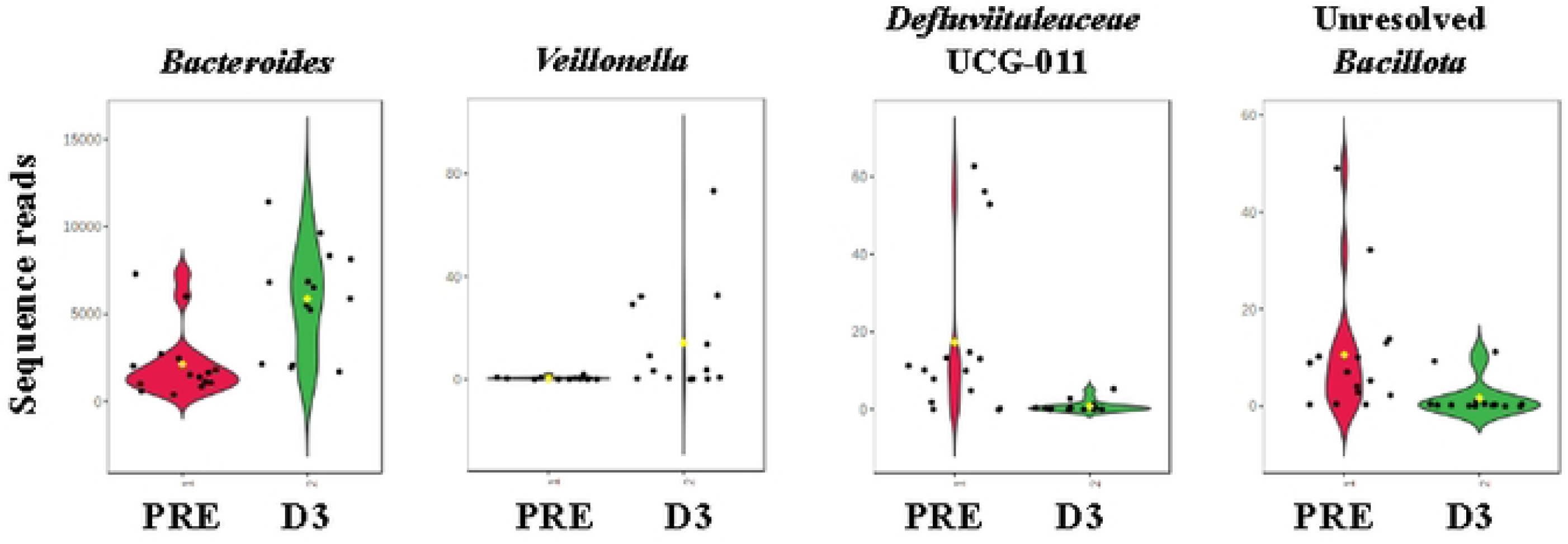
Violin plots comparing pre-colonoscopy and Day 3 for the four ASVs with the lowest *P*-values.

### Correlations between dietary intake and microbiome

The exploratory correlational analyses indicated that total energy (r=0.5815, *P*=0.0297), total carbohydrate intake (r=0.6860, *P*=0.0067), and total sugars (r=0.7754, *P*=0.0011) consumed on D2 post-colonoscopy were all significantly positively associated with the relative abundance of Christensenellaceae_R-7_group on D 3 post-colonoscopy (**Fig 7A-C**). These significant correlations were maintained after controlling for average pre-colonoscopy total energy intake. No other significant relationships were reported for any other dietary intakes and relative abundances of other ASVs at the different timepoints, though total energy intake during the pre-colonoscopy period was partially negatively associated with the relative abundance of Christensellanaceae_R-7_group at pre-colonoscopy (r=-0.4830, *P*=0.0682) (**Fig 7D**), and the consumption of sugars on D10 was also partially positively correlated with the relative abundance of Bacteroides on D11 (r=0.4867, *P*=0.0658) (**Fig 7E**).

**Fig 7.**
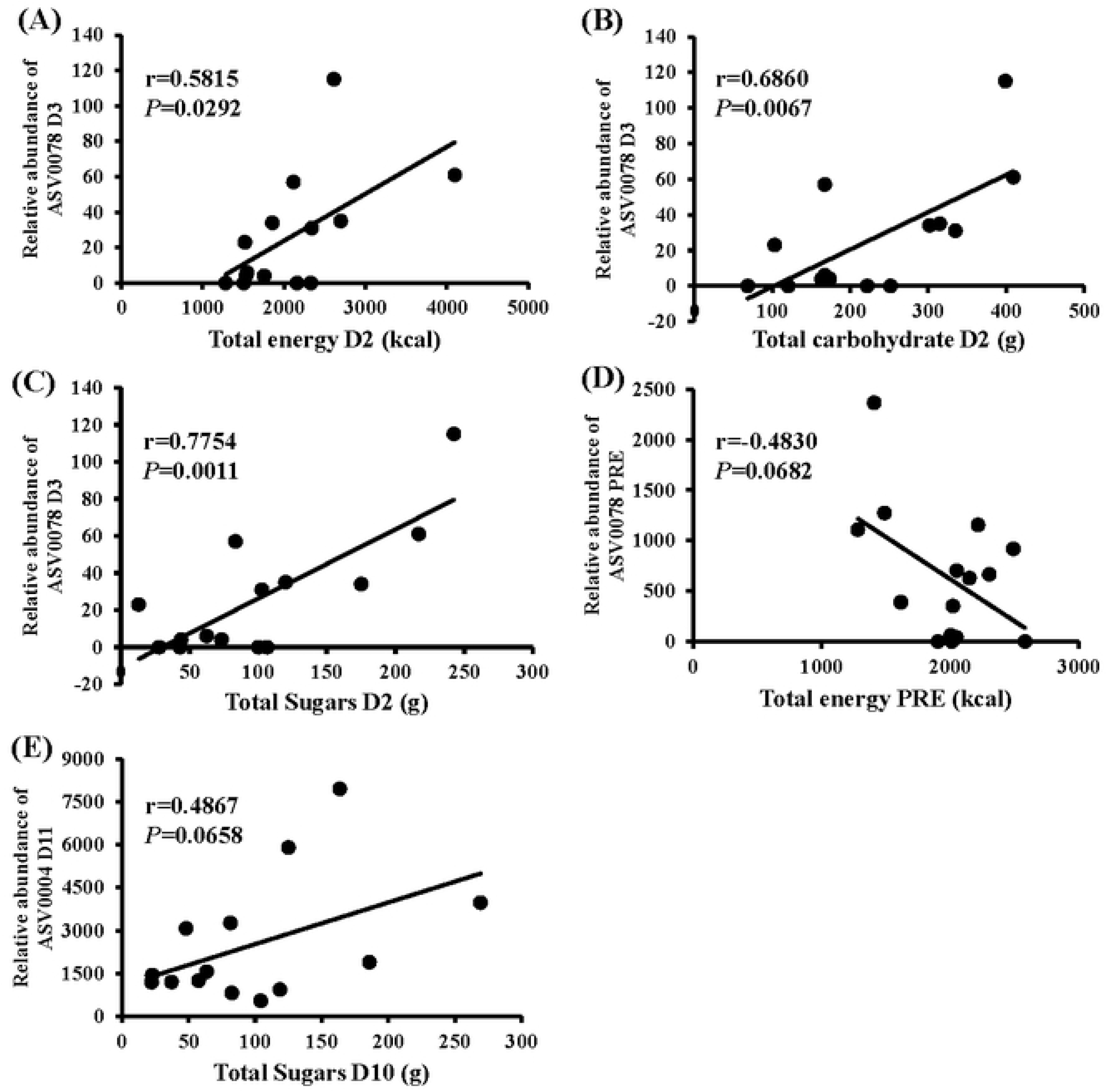
Relationships between dietary factors and the relative abundance of ASVs at multiple timepoints during the study. ASV0078 is Christensenellaceae_R-7_group, ASV004 is Bacteroides

## Discussion

The goal of this study was to determine the early pattern of microbial repopulation relative to a subject’s free-living dietary intake. In particular, we focused on dietary fiber intake because it is the strongest driver of microbial pattern and diversity(19, 20)]. Baseline fiber intake was higher than at Day 1 post-colonoscopy but thereafter, returned to baseline levels. Marked intersubject microbiome beta diversity was observed and three days post-colonoscopy, select taxa were depleted acutely (e.g., within the phylum Bacillota), while a few taxa were enriched compared to pre-colonoscopy levels (genus Parabacteroides, Bacteroides, Lachnoclostridium, Veillonella, Gamella, and Pasteurellaceae). These changes were lost by Day 5 when these ASVs were not significantly different from baseline at the level of the genus. Not surprisingly, marked inter-individual beta-diversity was observed among the study participants. Research has shown that the composition of the human gut microbiome varies considerably among individuals which can be explained by differences in host factors such as gender, age, ethnicity, environmental factors such as diet, lifestyle, and medication(21–25)] and also hereditary factors(26, 27)]. Here, significant differences were observed in the relative abundance of different ASVs in the fecal microbiome at Day 3. This result may be attributable to the short-term variability in the subject’s dietary intakes as has been reported by others(28)] and also as a result of the transient perturbation caused by the bowel cleansing associated with the colonoscopy procedure(5, 6)] leaving them energy deficient.

A few ASVs were identified as being differentially enriched 3 days post-colonoscopy including Gram-negative (Bacteroides, Parabacteroides, Pasteurellaceae) and organisms that are common but usually present at low levels when detected in feces (Veillonella, Gemella). The taxa that were depleted on Day 3 post-procedure included all the main species seen in feces associated with fiber digestion and the usual microbial functions (primarily of the phylum Bacillota). For example, bacteria in the genus Ruminococcus that are strictly anaerobic carbohydrate fermenters, have been studied in their animal hosts and found to serve the role of degrading complex polysaccharides into a host of nutrients for their host(29)]. Furthermore, bacteria from the genus Christensenellaceae _R-7 group belonging to the family Christensenellaceae have been reported to be negatively associated with visceral fat mass(30, 31)] and in mice, shown to be an indicative marker for lean phenotype(27)].

Exploratory correlational analyses were conducted to understand the relationships between dietary intake and changes in the microbial species. Individuals that consumed higher amounts of total energy, total carbohydrates, and total sugar on Day 2-post colonoscopy had greater relative abundances of this bacteria, the next day. Also, Roseburia, an anaerobic, gram-positive bacteria that was depleted post-colonoscopy is a butyrate-producing bacteria implicated in maintenance of energy homeostasis and prevention of intestinal inflammation(32)]. The decline in Bacillota observed Day 3 post-colonoscopy observed has also been reported in a recent study(6)]. For comparison, **Table 2** presents the results of past studies performing similar research. Mai and colleagues were the first to investigate the effects of bowel cleansing/colonoscopy on microbial composition to show that overall in 3 out of 5 subjects microbiota profiles after colonoscopy were similar to each other but different from pre-colonoscopy profiles(33)]. Further, Jalanka et al.(5)] and Smarr et al.(8)] demonstrated that colonoscopy-associated bowel cleansing decreased the microbial load by 31-fold and alpha diversity respectively, and they assessed active repopulation 2 weeks later. Similarly, Nagata et al. demonstrated that compared to a no-bowel prep group (controls), microbiota composition after bowel prep was significantly reduced immediately after the prep but not at 14 days after the procedure(7)]. Very recently, Chen and colleagues also demonstrated that the relative abundance of Firmicutes was reduced by 30% 3 days post-colonoscopy but returned to baseline levels by day 7 after subjects were supplemented with the probiotic *Clostridium butyricum* for 60 days following the colonoscopy procedure(6)]. Altogether, these findings are indicative of the drastic effect of the bowel cleansing in depleting beneficial microbiome and suggest that at least 2 days of food intake would be needed to repopulate the gut microbiome after this disruption. Others have even suggested that the bowel cleansing associated with colonoscopy has long-lasting effects on the homeostasis of the gut microbiota, particularly resulting in a decrease in the abundance of the beneficial bacterial population, Lactobacillaceae(34)].

**Table 2.** Summary of studies investigating the microbiome after a bowel cleanse.

| Author / publication year / PMID | Sample size / population | Intervention and duration of data collection | Primary findings |
| --- | --- | --- | --- |
| Mai et al./2006/ PMC1856456 | N=5 subjects underdoing screening colonoscopy | Fecal microbiota analyzed by denaturing gradient gel electrophoresis. | In 3 of 5 patients, the two profiles detected after the colonoscopy were more similar to each other and clearly different from the pre-colonoscopy profile; in the other 2 patients, there was less of a difference between the pre- and post-colonoscopy profiles. |
| Jalanka et al./2015/ PMID: 25527456 | N=23 healthy subjects from Nottingham University | Group I - split dose of 2 L of PEG solution; 1<sup>st</sup> liter in the evening and the 2<sup>nd</sup> liter the | Total microbial load was decreased by 31-fold and 22% of participants lost the subject-specificity of their microbiota. Bacterial levels and |
|  | hospital assigned to two groups | following morning. Group II- single dose of 2 L of PEG solution on the morning of the study. Fecal samples collected at baseline (a day before the bowel cleanse), immediately after the lavage, and on days 14 and 28. Microbiota composition was analyzed by phylogenetic microarray (HITChip). | community composition were restored within 14 days, with the rate of recovery being dose dependent: single dose PEG had a more severe effect on the microbiota composition than double dose, and importantly, increased the levels of Proteobacteria, Fusobacteria and bacteria related to Dorea formicigenerans. |
| Smarr at al./2017/ PMID: 29582916 | N=1, age 67 y | Larry Smarr quantified daily shifts in his gut microbiome composition from July 1-31 (colonoscopy was on July 7) using 16s rRNA gene sequencing. Colon cleanse was done with PEG solution. | Alpha diversity (average phylogenetic score) dropped from an average of 34-39 in the days prior to an average of 22-24 in the two days immediately post colonoscopy and returned to baseline levels in ~9 days. PCoA plot indicated that pre-colonoscopy samples clustered together with samples collected two weeks after colonoscopy, while samples collected immediately post colonoscopy to about day 9 after were characterized by large movements in PCoA space. Also, genus Rosburia jumped over 9-fold post-colonoscopy, and genus Coprococcus dropped by 40-fold post-colonoscopy. |
| Nataga et al./ 2019/PMID: 30858400 | Bowel prep group (n=8) and no-procedure group brought from literature (n=23). Study population: Tokyo, Japan. | In the bowel prep group fecal samples collected before, during (immediately after the bowel prep), and 14 days after bowel prep. In control group, fecal samples collected on Day 0 and Day 14. 16S rRNA sequencing | Compared with controls, microbiota composition was significantly reduced immediately after the prep but not at 14 days afterward. |
| Chen Zhenhui et al./ 2024/ PMID: 38429821 | N=11 that underwent colonoscopy (7 in intervention, 4 in control group). Chinese population. | Sixty days study. N=7 supplemented with 3 pills of *C. butyricum* immediately after colonoscopy for 20 days. N=4 no supplementation after | *Firmicutes* and *Bacteroidetes* changed on day 0 and on day 1 and 3 after colonoscopy ($P<0.05$). Almost all bacterial phyla were at their highest or lowest abundances on day 3 and *Firmicutes* decreased by 30% ($P<0.05$). The *Firmicutes* to *Bacteroidetes* ratio decreased during |
|  |  | colonoscopy. Fecal samples were collected before bowel lavage(40), during colonoscopy (day 0) and after colonoscopy (day 1, 3, 7, 14, 30, and 60). 16S rRNA gene sequencing | colonoscopy and on the third day after colonoscopy then returned to baseline levels on day 7. *Firmicutes* decreased by 16.1% in *C. butyricum*-treated subjects on three days after colonoscopy compared with 27.9% in the control, while *Bacteroidetes* increased by 13.4% on three days after colonoscopy compared with 22.4% in controls. |
| Drago et al./2016/<br>PMID: 27015015 | 10 patients | Fecal sample collected immediately after a 4 L PEG-based (SELG Esse) bowel lavage for colonoscopy and 1 month thereafter. 16S rDNA Ion Torrent profiling of fecal samples | Significant decrease in *Firmicutes* abundance and an increase in Proteobacteria abundance immediately after the colon cleansing and 1month after. Significant reduction in family *Lactobacillaceae* and an increase in *Enterobacteriaceae* abundance were observed immediately after the colonoscopy, whereas 1 month after, these families were significantly lower compared with pre-colonoscopy samples. *Streptococcaceae* were increased 4-fold 1 month after the colonoscopy compared with pre-colonoscopy samples. |

Summaries of literature derived from a PubMed search on the topic of microbiome changes after a colonoscopy. PEG, Polyethylene glycol.

### Influence of food intake

In the present study, fecal samples taken on the morning of Day 3 post-colonoscopy would be expected to be more reflective of the food consumption pattern immediately following the colonoscopy procedure. Total fiber intake on the day of the colonoscopy declined by about 14% in comparison to pre-colonoscopy intake and came back to pre-colonoscopy levels thereafter. It is acknowledged that on the day before the procedure, subject food intake was very low as a result of the colonoscopy preparation, and we believe that from Day 0 – Day 5, early food intake peaks were a result of “catch up” food intake after no energy intake during the preparation period. The subjects having limited energy intake for over 24 hours followed by undergoing a colonoscopy with an anesthetic would cause a period of physical stress and increase energy demands. Thus, both the composition, quantity, and timing of food intake will influence the process of repopulation.

In our study and in previous studies that have explored the role of microbiome changes following bowel cleanse have primarily utilized PEG as the bowel lavage solution. In real life clinical practice this is not always the case. Several endoscopists choose to use alternative solutions like magnesium citrate and sodium picosulfate. Some providers may also choose to use a combination of these laxatives and have extended periods of dietary restrictions prior to the colonoscopy, like a low-fiber diet for up to one week before the procedure. The effect of these solutions on the gut microbiome has not been studied nor has it been compared to the changes observed after the use of PEG. In an animal study where PEG and citrate-based laxatives were fed to mice, both laxatives were found to have an effect on changing their gut microbial composition with a highPEG dose reducing alpha and beta diversity(35)].

### Strengths and limitations

A major strength of this study is that it reflects real-life repopulation of the microbiome after a screening colonoscopy preparation in an outpatient setting. Another strength is the repeated data collection at baseline and over two weeks post-colonoscopy. Subjects were 100% compliant in providing food records and fecal samples and this increased the statistical power to detect differences over time. As described above, other investigators have studied microbiome changes associated with the colonoscopy procedure and shown that the bowel cleansing procedure significantly alters microbiome profiles(5, 8, 34, 36)]. This study expands on those data by collecting detailed dietary intake information concurrently with studying microbiome changes. Diet is a very important predictor of microbiome response, but this relationship is complex; thus, understanding inter-individual dietary intake patterns during the important period following the colonoscopy procedure is important in informing dietary intervention trials aimed at promoting gut health. Limitations of this project included the relatively small sample size and the design being a single-arm study. Future studies should collect data from a control group, over the same duration in individuals who did not undergo a colonoscopy. Similar studies conducted among subjects with medical illnesses like inflammatory bowel disease, irritable bowel syndrome, and other illnesses where gut microbial dysbiosis has been established would be highly valuable (37)]. Further, the effect of the subject consuming no food for 24 hours could also be controlled for. In analyzing these data, we considered the effect of prep-day starvation on subsequent food intake - anticipating that the total energy consumed on Day 1 post-colonoscopy might be higher to make up for low energy intake the day before the procedure. We observed that this energy deficit took more than one day to accommodate. Given common comments by patients, we also suspected that on Day 1, subjects would choose high-fat or high-sugar foods as a reward for completing their preventative health screenings and particularly, for complying with the fairly laborious prep. We did not find this to be the case. Lastly, food intake is known to be different between weekdays and weekend days (38, 39)]. We analyzed the timing of the colonoscopy relative to the weekend, and no clear pattern was found to weekend eating as affecting the results. To control for this potential effect, future investigations should study subjects who have all had their colonoscopy procedures on the same day of the week.

## Conclusion

These findings demonstrate the magnitude of the significant decline in microbial relative abundance and diversity immediately post colonoscopy. The timing of repopulation aligned with changes in fiber intake post-procedure. These data highlight the importance of nutrition after a screening colonoscopy in reestablishing a healthy microbiome. Future investigations should determine whether there is benefit, to advocating that post-colonoscopy patients increase their fiber intake to support the repopulation of the microbiome and resumption of gut health.

## Author contributions

Ghouri YA- data analysis, writing and clinical perspective; Ericsson AC- data generation and analysis, manuscript writing; Anderson JM- recruitment and screening, dietary data lead and clinical trial management; George JG- assisted with recruitment, subject coordination and data generation; Parks EJ- study design, data analysis, and manuscript writing; Anguah KOB: study design, microbiome analysis, scientific data interpretation, manuscript writing.

